# A Modular, Adaptive, Deep-Learning-Based Brain-VR Interface

**DOI:** 10.1101/2022.11.10.515931

**Authors:** Maryna Kapitonova, Zacharias Häringer, Eric Bongk, Tonio Ball

**Affiliations:** Department of Neurosurgery, IMBIT, University of Freiburg, Germany

**Keywords:** brain, neurocontrollers, artificial intelligence

## Abstract

Brain-Computer Interfaces (BCIs) may open up new possibilities for Virtual Reality (VR) applications: BCIs may be used for active brain control of VR avatars, or to make VR content passively-adaptive based on information decoded from ongoing brain activity. Application domains for such Brain-VR Interfaces (BVRI) include medical and healthcare, entertainment, and education. Conversely, VR technology also opens up new possibilities for BCI research and development: E.g., gamified immersive BCI paradigms may improve subject engagement and long-term motivation, helping to study learning and adaptivity in the BCI-control context. Previously, we have demonstrated a first adaptive, deep-learning-based online BCI for the control of robotic assistants. Here, we describe the extension of this setup to a modular, extensible, VR-compatible online BCI setup. We describe how we integrated a classical active BCI control paradigm using motor imagery into a gamified interactive VR scenario, designed to enhance the long-term motivation of subjects. We also present an initial quality assessment of electroencephalographic (EEG) signals acquired with a dry-electrode system. We anticipate that the presented modular adaptive Brain-VR Interface will help to understand and facilitate (co-)adaptivity during long-term BCI usage.

## I. INTRODUCTION

Virtual Reality (VR) technology has matured to provide highly immersive VR experiences using customer-grade commercial VR systems. VR applications may be used for rehabilitation in neurologic patients, have been widely adopted by the gaming industry, and open up many new possibilities in education and training [5]. In the form of the metaverse, VR may have a substantial impact on the future of work. However, already before VR became a mainstream technology, it was noted that from the integration of VR and BCI technologies particularly interesting and unique applications may emerge.

Previously we have developed an adaptive deep-learning BCI for robot control in (non-virtual) reality. As the reliable classification of brain signals related to directional commands (e.g., simply thinking “go right” etc.) is challenging to achieve with currently available non-invasive brain signal decoding techniques, in our previous work we used a hybrid deep ConvNet approach to infer multiple surrogate mental tasks from non-invasive EEG, using 40-Hz low-pass filtered EEG data to decode 5 different mental tasks. This shared-autonomy BCI concept allowed reliable control of complex sequential actions of the robot assistant. (Re-)analysis of the data from these experiments showed that adaptivity on the deep decoder side was a crucial factor for success; however, at least so far we did not find clear evidence for adaptivity on the level of the brain signals that subject produced from session to session. One possible explanation for this lack of detectable learning and adaptivity on the subject level may be that training over longer time scales (weeks to months) would be required.

In our previous work as described above, we used wet (gel-filled) EEG electrodes [4]. Dry electrode systems may be more convenient to use, and thus especially helpful for long-term studies, but can pose challenges with respect to signal quality. Previously we studied the signal quality of EEG when using two commercial head-mounted displays (HMDs), the Oculus Rift and the HTC Vive Pro, on the quality of 64-channel wet-electrode EEG measurements. We found that expectedly, the HMDs consistently introduced artifacts, especially at the line hum of 50 Hz and the HMD refresh rate of these specific models (90 Hz), as well as their harmonics. However, as “good news” for EEG-BCI-VR studies, the frequency range that is typically relevant for non-invasive EEG BCIs below 50 Hz, was unaffected by strong artefacts. One aim of the present study was to investigate dry EEG signal quality during HMD usage.

Most of the studies for EEG-controlled VR environments are motivated from the neurogaming perspective [6], and it is not always entirely clear to which extent such applications utilize brain signals and not artefacts. Here, we aim to show the feasibility of portable BCI-VR set-ups for designing experiments, aiming to answer scientific questions from the neuroscience side, as well as from the algorithm development perspective. To enable long-term BCI-VR studies, here we describe how we have extended our adaptive deep-learning-based BCI setup to gamified VR applications. In the present work, we were also particularly interested in the signal quality of dry electrode recordings while wearing a HMD. Previously, Yao and colleagues demonstrated the feasibility of SSVEP-based online BCI applications in VR environments, but without using deep learning on the decoding side and utilizing wet EEG electrodes [3]. To our knowledge, the presented here system is the first modular online VR-BCI utilizing dry-electrode recordings and powered by adaptive deep learning.

## II. BCI-VR Set-up

Throughout the design of our BCI-VR system, we followed a modular approach. Our set-up integrates VR components, EEG acquisition devices, and decoders in an extensible and exchangeable fashion: The VR application itself can be exchanged, data acquisition does not depend on the specific EEG hardware, and decoders can be easily exchanged and/or customized for different paradigms as well as channel counts and electrode layouts. In addition, the set-up can be extended for the acquisition of other multimodal biosignals, such as the galvanic skin response, heartbeat, eye movements, etc., if it is required by the experimental paradigm. The schematic of our set-up is shown in Fig. 1.

**Fig. 1.**
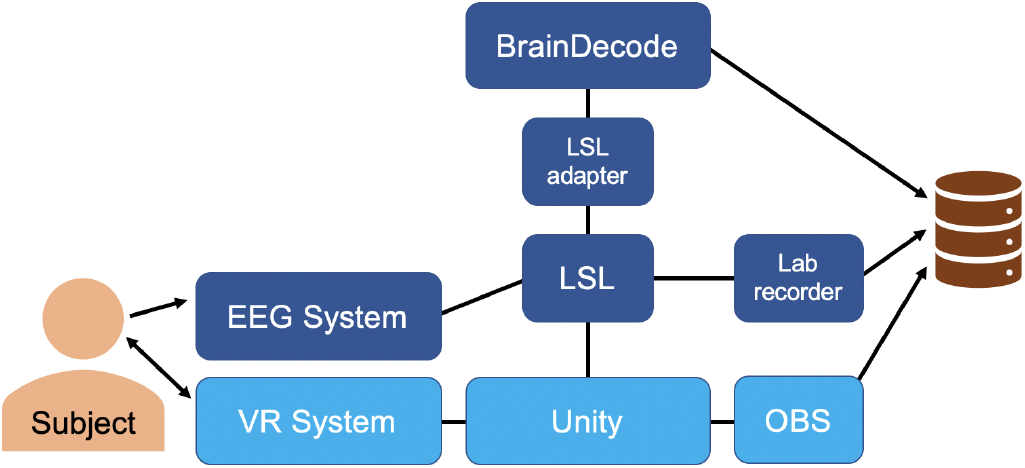
Schematic of our modular BCI-VR set-up.

### A. Data Acquisition

EEG signals were acquired using portable eego sports 64 channels amplifier produced (ANT Neuro). The Waveguard Touch dry EEG cap has 64 coated Ag/AgCl flexible polymer electrodes placed according to an equidistant hexagonal layout. For motor tasks, as we are mainly interested in *α* and *β* frequency bands, we set the sampling rate to 500Hz.

### B. Unity Game Engine

We used the cross-platform game engine Unity developed by Unity Technologies to develop our game. The threedimensional (3D) version, also known as Unity3D offers a broad component system, including audio systems, physic systems, customizable rendering pipelines and a scripting API for the C# programming language. Recently Unity was adopted by researchers for experimental design, offering a broad range of possibilities for behavioral studies [8]. In addition, the package system allows to use LSL4Unity^1^ for synchronization with other devices and support for the integration of various VR-systems.

### C. Communication via LSL and EEG Preprocessing

We relied on Lab Streaming Layer (LSL) to synchronize the devices within the set-up and to record the data, i.e., EEG data, events generated within the Unity VR game, Braindecode-online output (predictions) etc. LSL is also involved in the preprocessing, filtering and downsampling the data to prepare it for deep learning decoder. In the current version 400 data samples are gathered for better processing speed and filter quality. Then, a 50-Hz notch filter with a quality factor of 30 was used to eliminate the power line hum, followed by a low-pass filter (40 Hz, 6th order Butterworth filter). Further, the signal was downsampled to 250Hz, which is adequate given 2s as the receptive field size and the frequency range of the most informative components. As a last step, we perform exponential moving standartization to normalized and center the signal around 0.

### D. BrainDecode-online

For a decoding module we used the online version of the BrainDecode toolbox [1], [2], containing popular architectures for EEG decoding, such as EEGNet, Deep4, ShallowNet, TCN and TIDNet. The module contains pretrained networks and provides a possibility to use them for online inference, as well as to fine-tune them online for a particular task or subject.

Once the decoder has gathered 600 samples, necessary for the network operation, it will start decoding the signal. After this a new prediction is made every 125 samples. The outputs of the decoder are accumulated in a ring buffer of size 6. The predictions stored in the ring buffer are then aggregated (we are currently evaluating different aggregation functions) and a final decoding output is generated and forwarded via LSL to Unity for informing the gameplay.

## III. Game Design

For user engagement and to ensure a rich experimental paradigm we designed a sci-fi VR game. The main purpose of the game is to facilitate longitudinal data acquisition, naturally inducing constraints on the game design. The objective of the game is to catch robots by performing motor imagery, thus collecting points. The game loop consists of short sessions (5 min) to collect as many points as possible, particularly good results will be saved on a leader board to spur the ambition of the player and to improve the replayability of the game. The obtained points additionally allow the user to unlock new environments; less competitive users can find motivation in exploring the three different levels (a space station, a laboratory at a lava lake and a planet surface with crystal formations), organised in a star structure. In-game locomotion is implemented using teleportation to reduce motion sickness.

The game has a *Training Mode* for collection of the training data, used to train a first subject-specific model, and a *Gaming Mode* with online control based on this initial model and allowing for online adaptation. Furthermore, with the purpose of debugging the game we implemented a *Replay Mode*, which allows to simulate the online EEG control.

## IV. Acquisition of training data

Participants were seated in an electrically shielded room. For the EEG measurement, the 64 passive electrode EEG cap as described above was used. The reference electrode was placed on the right mastoid and the ground electrode on the left. The EEG was recorded via the eego software on a Laptop and streamed via the software plugin to LSL. The VR headset (Valve Index), was placed on top of the EEG cap. To interact with the VR environment, the participant used a Valve index controller with each hand. The VR headset was controlled via the Unity engine and sent game-related event signals and the X-, Y-, and Z-location of the controllers to LSL. All the data are timestamped by LSL and stored on connected HD.

After starting the VR game in Training Mode, the participant has to move to the training area (Fig. 3B). The player can now press the A-Button of the controller to spawn a robot. There are two different kinds of robots, which we call Robot Smiley; with a round head, and Robot Buddy; with a square head. Both robot types can change color. In the nonactive state, both robots’ color is grey. If the player faces the robot and looks directly at it, the robot changes its color to yellow, indicating that the participant is supposed to sit still and not move. This stage is called the *resting phase*. Here, the participant should refrain from movement for 5 s. If the timer runs out, the robot is going to disappear.

**Fig. 2.**
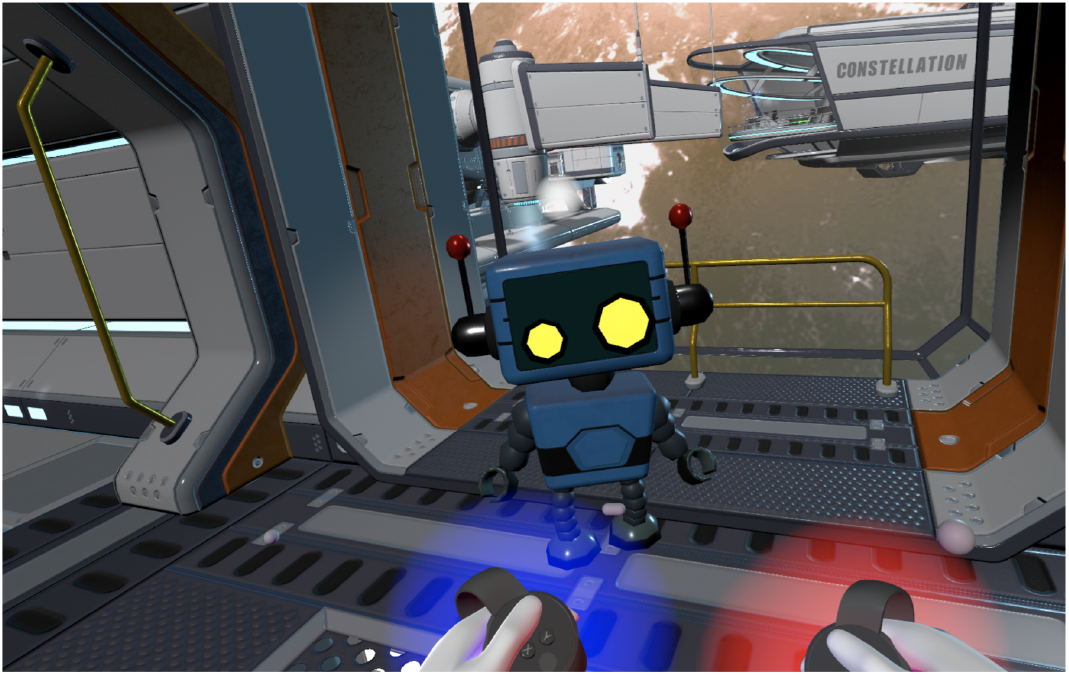
Game screenshot example. In the online mode, red and blue glow around the hands is controlled by imagined right and left hand movements. Color of glow must match color of robots to collect them.

**Fig. 3.**
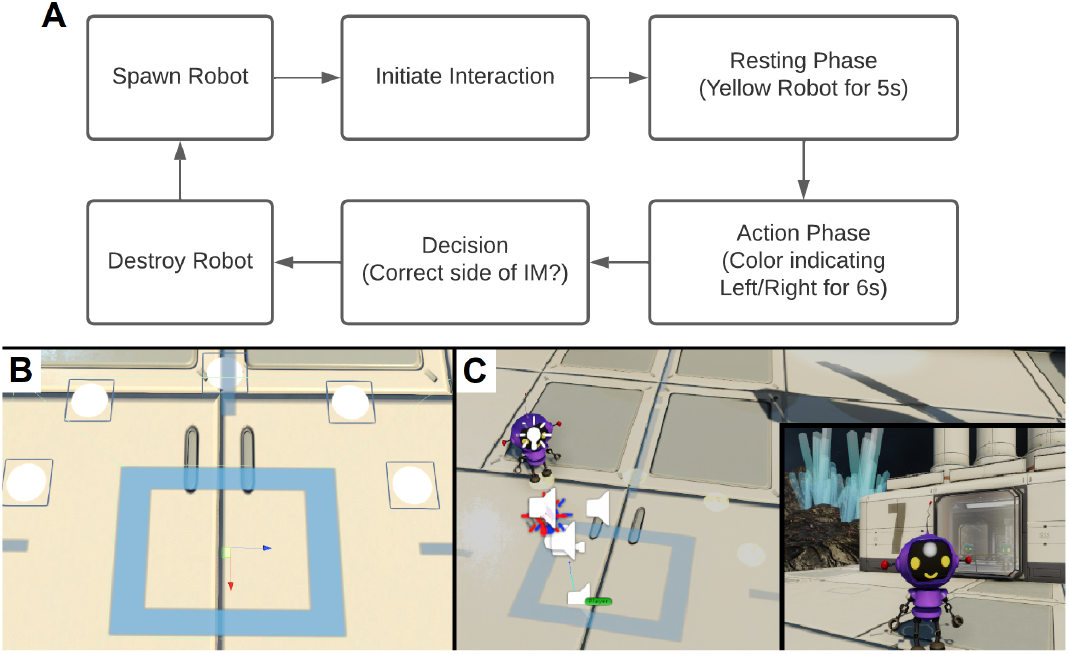
A) Interaction overview diagram. B) Top-down view of the training area (blue square). Entering the area triggers the appearance of five possible spawn locations for the robots (white circles). C) Unity editor view on an interaction between player and robot (left), and from the point of view of the participant (right)

After the game detects that the player is not moving, the action phase starts. The robot changes its color randomly to red, purple, green, or blue (Fig. 4). Depending on the color, the player has to imagine the movement of the right or left hand. Red and purple stand for the right side and blue and green for the left. This *action phase* lasts for 6 s. At the end of the action phase, the robot displays a short animation indicating success or failure. The game sends event information about robot type, robot color, robot location, interaction outcome, and the associated time point. All steps of this interaction cycle are visualised in Fig. 3A.

**Fig. 4.**
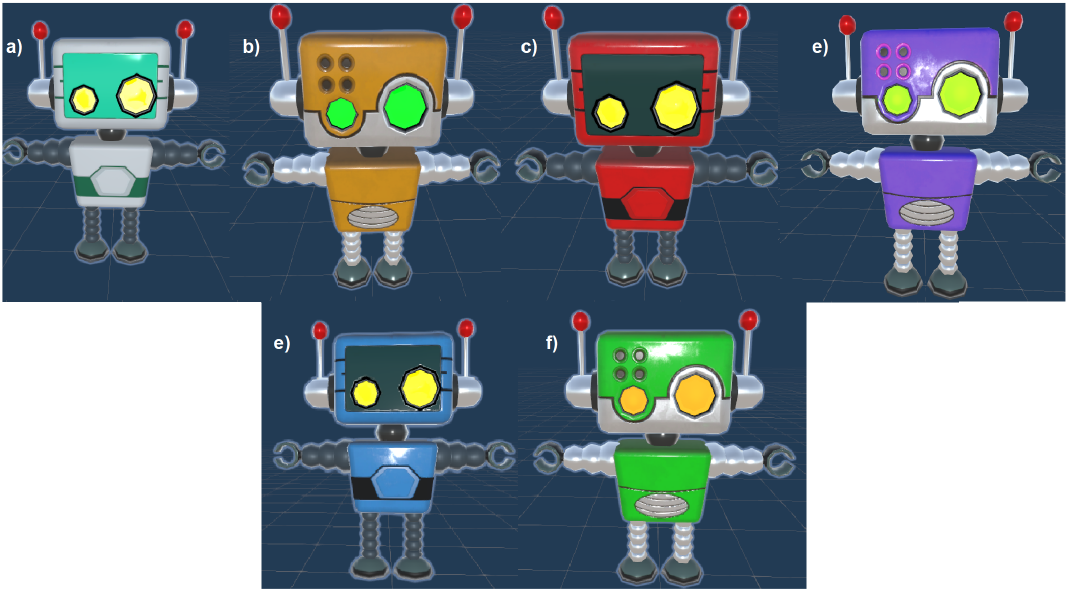
Robot “Buddy” color states. These Unity models were bought on the Unity store from the user Redhead Robot and slightly edited. a) robot outside of encounter. b) encounter is triggered participant is supposed to get into a resting state. c) and d) are both colors to be associated with right hand movement. e) and f) are associated with left hand movement

**Fig. 5.**
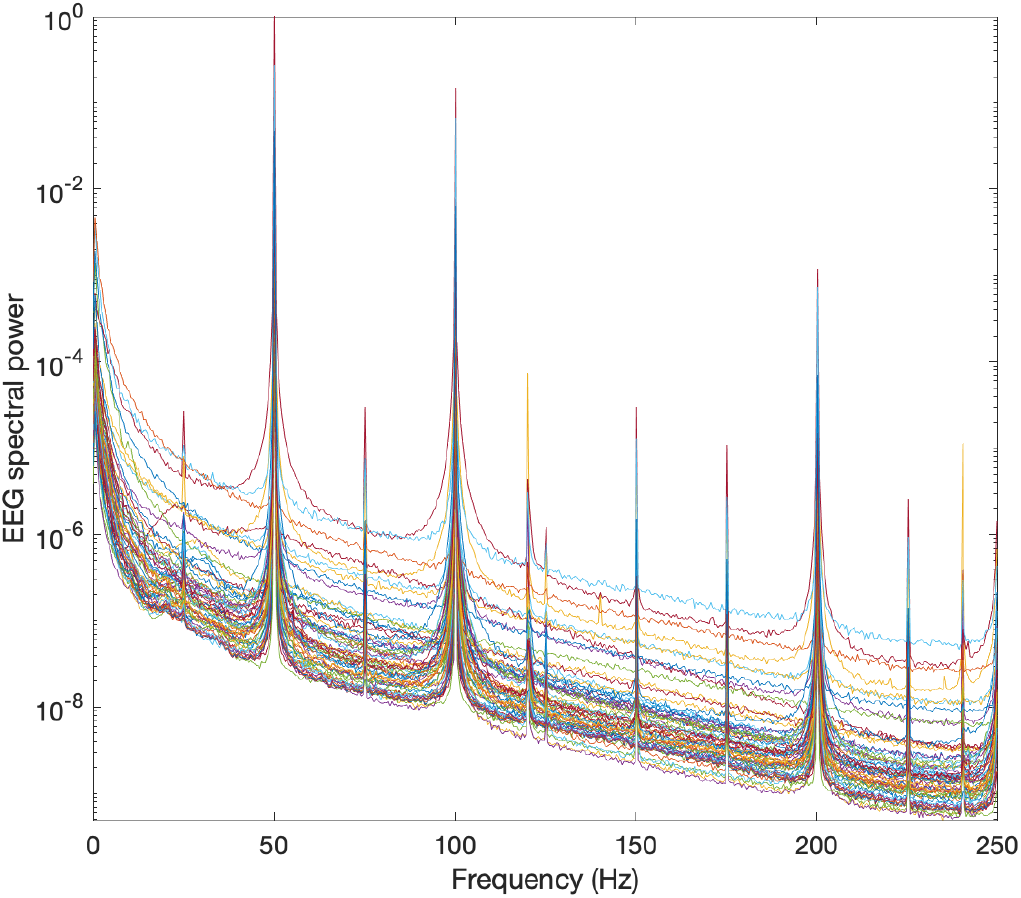
Absolute EEG spectral power for all 64 channels. Expected artifacts at line hum frequencies and HMD refresh rate (120 Hz).

**Fig. 6.**
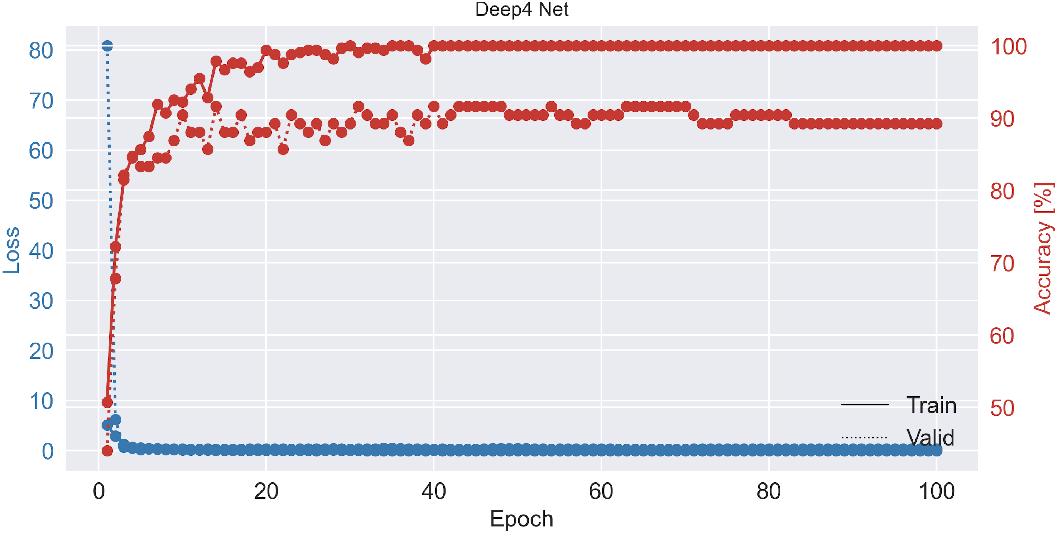
Decoding results for right vs. left hand movement using the Deep4 architecture.

## V. Dry EEG Quality and Decoding Results

To evaluate the signal quality of the dry electrode recordings during HMD usage, we analyzed repetitive right and left hand movements in time and frequency domains, as well as with respect to decodability.

We calculated movement-related potentials (MRPs), movement-related spectral power changes, as well as absolute spectral power of ongoing recordings at all channel positions. We decoded the two movement classes be training the Deep4 model on an 80:20 split of the data.

We found MRPs and movement-related spectral power changes in the expected frequency ranges up to 40 Hz, with suppression in the *μ* and *β* bands as well as a post-movement power rebound. Absolute spectra showed the expected artifacts at the line hum frequency and harmonics, and at the HMD refresh rate (120 Hz). Decoding accuracy of right vs. left hand movement trials using the Deep4 model was 89.3%.

## VI. Conclusions

In conclusion, the proposed brain-VR interfacing set-up is:

- provides gamification to experimental design increasing subject engagements and return rate.
- provides a testbed for prototyping expensive experiments in the field of BCI-controlled robotics (e.g., drone control, swarm control),
- has a potential to help bringing BCIs out of the lab, with VR as an intermediate step, and given the quick set-up time for dry EEG system and portability of amplifier. In this context, future directions also include implementing decoders on edge devices as well as integrating cloud-based functionality, to even more enhance the portability of the set-up.
- might have a potential to mitigate “BCI illiteracy”, as it was shown already that utilizing DL decoders leads to better performance in users, which are struggling to learn BCI control [7],
- a step to unified experimental design, allowing to collect within one paradigm data suitable for multiple decoding tasks and scientific questions, which in turns increases the effectiveness of resource utilization,

We anticipate that modular, adaptive, deep-learning-based brain-VR interfacing technology, as described here, will develop into a versatile research tool, and help paving the way for a broad spectrum of novel BCI-VR applications, opening up new possibilities of how the human brain may interact with immersive virtual environments.

## Acknowledgment

We would like to thank Henrik Bonsmann and Xi Wang for their help. Conflict of interest statement: TB and MK hold shares of the NeuroMentum AI GmbH, which develops BCI concepts and technology.

Copyright by IEEE.

## Footnotes

1 https://github.com/labstreaminglayer/LSL4Unity

